# Complex regulation of gamma-hemolysin expression impacts *S. aureus* virulence

**DOI:** 10.1101/2022.10.19.512982

**Authors:** Mariane Pivard, Isabelle Caldelari, Virginie Brun, Delphine Croisier, Michel Jaquinod, Nelson Anzala, Benoît Gilquin, Chloé Teixeira, Yvonne Benito, Florence Couzon, Pascale Romby, Karen Moreau, François Vandenesch

## Abstract

*Staphylococcus aureus* gamma-hemolysin CB (HlgCB) is a core-genome encoded pore-forming toxin that targets the C5a receptor, similarly as the phage-encoded Panton-Valentine Leucocidin. Absolute quantification by mass spectrometry of HlgCB in 39 community-acquired pneumonia (CAP) isolates showed considerable variations in HlgC and HlgB yields between isolates. Interestingly, when testing the hypothesis that HlgCB might be associated with severe *S. aureus* CAP, we found that a high level of HlgCB synthesis was associated with mortality in a rabbit model of pneumonia. To decipher the molecular basis for the variation in *hlg*CB and *hlg*B expression and protein production among strains, different regulation levels were analyzed in representative clinical isolates and reference strains. Although HlgC and HlgB are encoded on a single operon, their levels were dissociated in 10% of the clinical strains studied. HlgCB amount and HlgC/HlgB ratio were found to both depend on promotor activity, mRNA stability and translatability, and on the presence of an individual *hlg*B mRNA processed from the *hlg*CB transcript. Strikingly, toe-printing and *in vitro* translation assays revealed that a single SNP in the 5’-UTR of *hlg*CB mRNA strongly impaired *hlg*C translation in the USA300 strain, leading to a strong decrease in HlgC but not in HlgB; the level of HlgB is likely to have been maintained by the presence of the processed *hlg*B mRNA. This work illustrates the complexity of virulence factor expression in clinical strains and demonstrates a butterfly effect, where subtle genomic variations have a major impact on phenotype and virulence.

**Author Summary:** The Gram-positive bacterium *Staphylococcus aureus* can provoke a wide range of infections due to its ability to produce a large diversity of virulence factors, including immune evasion molecules, adhesins, and toxins. Some of these toxin-encoding genes are localized in mobile genetic elements, and are thus not present in all strains, whilst others are encoded in the core-genome and present in all strains. Gamma-hemolysin CB is a core-genome encoded toxin but its amount varies between community-acquired pneumonia isolates. The regulation mechanisms underlying this variation however, are not well characterized. Here, we show that gamma-hemolysin expression levels vary largely among clinical strains and that, when highly produced, it induces high mortality in a rabbit model of pneumonia. The molecular basis for the variation in gamma-hemolysin expression depends on multiple mechanisms including promoter strength, transcript stability and processing, and translatability (i.e. the amount of protein that is synthetized by the ribosome for a given transcript). Incredibly, all these factors rely on a subtle genetic modification. This work emphasizes the importance of the disparity in virulence factor expression among clinical isolates and points the extreme complexity of the molecular mechanisms underlying their regulation, rendering the prediction of virulence for a clinical isolate difficult.

## Introduction

The Gram-positive bacterium *Staphylococcus aureus* can provoke various severe diseases in humans such as bacteremia, endocarditis, and pneumonia. Its capacity to develop a wide range of infections is due to its ability to produce a large diversity of virulence factors including immune evasion molecules, adhesins, and toxins (1,2). *S. aureus* accounts for approximately 25% of nosocomial pneumonia (3) and can be responsible for severe community-acquired pneumonia (CAP) with high mortality rates (4–7). In this last context, the bi-component pore-forming toxin (PFT) Panton-Valentine leucocidin (PVL), which targets the C5a and C5L2 receptors on neutrophils, monocytes, and macrophages, and the CD45 receptor on leukocytes (8,9), has been associated with severity independently from other virulence determinants and methicillin resistance (4). Nevertheless, although PVL is present in almost 50% of strains isolated from patients with severe CAP (4), the remaining cases are caused by PVL-negative strains and are still associated with 30% mortality (4), suggesting that other virulence factors are involved. One plausible candidate is gamma-hemolysin CB (HlgCB), which is present in almost all strains (10) and also targets the C5a receptors of myeloid cells (11,12). Its genes are part of the gamma-hemolysin *hlg*ACB locus; *hlg*A is upstream the *hlg*CB operon (Fig. 1A) (13), allowing the formation of two toxins, HlgAB and HlgCB. Although HlgAB is known to be involved in bloodstream (14) and corneal (15) infections by binding the CCR2 and CXCR1/2 receptors on neutrophils, macrophages, and monocytes, and the DARC receptor on erythrocytes, HlgCB is not typically associated with any specific disease. Recently, the semi-quantitative proteomic analysis of exotoxins applied to 136 strains isolated from severe CAP suggested that high HlgCB production in PVL-negative strains could be associated with higher mortality. It also showed that HlgB and HlgC levels were not correlated, contrarily to other operon-encoded virulence factors such as LukDE and LukSF-PV (16). We thus decided to investigate the molecular mechanisms leading to HlgC and HlgB expression variations among clinical isolates and assess whether HlgCB is a major determinant for pneumonia severity in a rabbit model (17). Besides identifying a new translatable *hlg*B mRNA as a maturation product of *hlg*CB, the present data shows that the variation in HlgC and HlgB production results from at least three parameters: promoter strength, *hlg*CB mRNA stability, and translation efficiency, which is dependent on a single nucleotide polymorphism (SNP) in the 5’UTR of *hlg*C mRNA. Furthermore, the high expression of HlgCB by clinical isolates induced high mortality in the rabbit model, a result in line with the hypothesis that HlgCB might be associated with severe *S. aureus* CAP.

**Figure 1.**
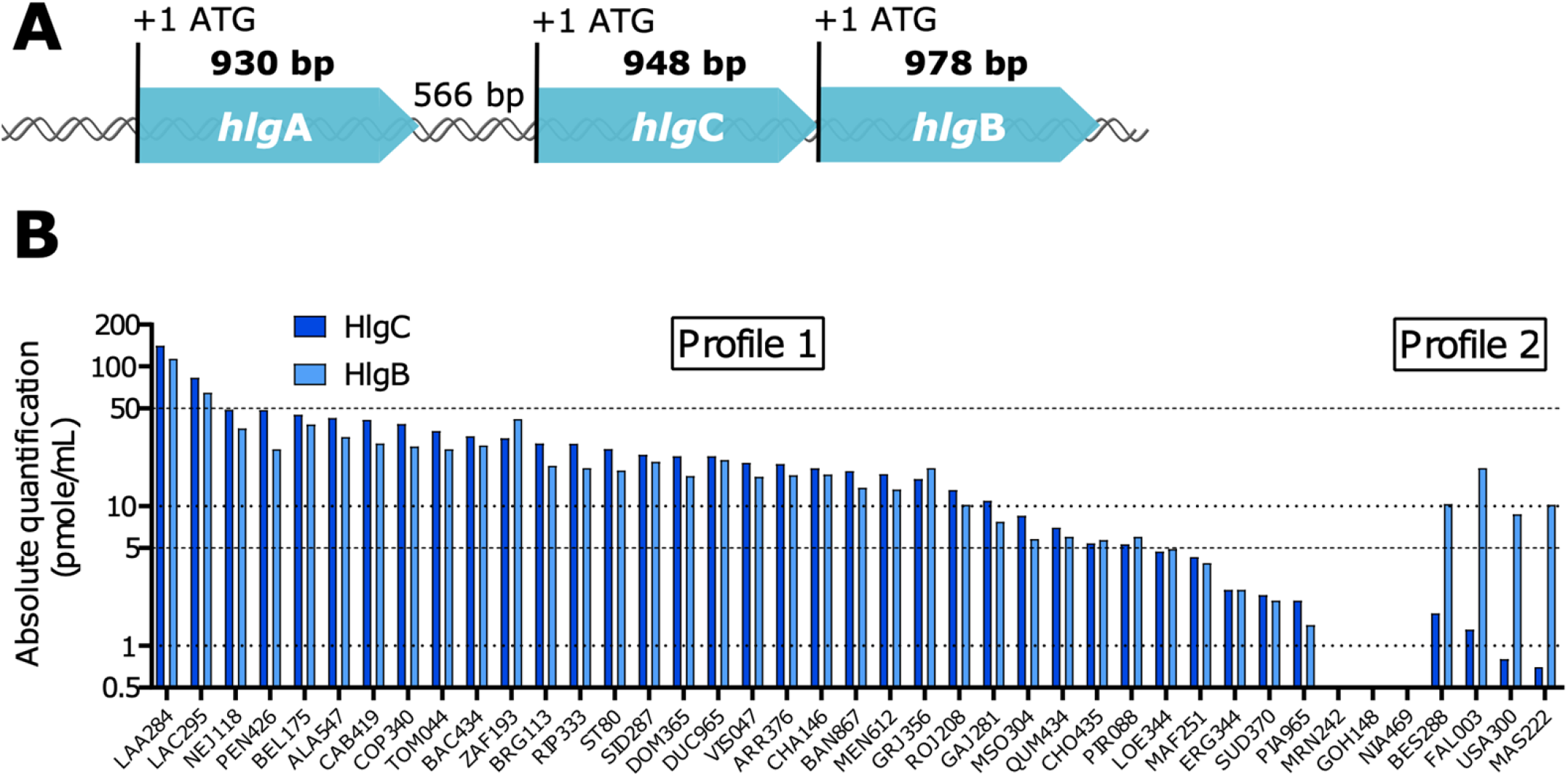
Gamma-hemolysin genomic locus: two distinct profiles and high variation in HlgCB production between clinical isolates. (A) Schematic representation of the gamma-hemolysin genomic locus. The size of the coding and intergenic regions are expressed in base pair (bp). (B) HlgC (dark blue) and HlgB (light blue) were quantified using isotope-dilution targeted proteomics on *S. aureus* culture supernatants. ST80, USA300, and 39 clinical isolates from severe-CAP were cultured in CCY for 8 h, supernatants were collected and analyzed by targeted mass spectrometry. Quantifications are represented on a log_10_ scale in pmole/mL. Two distinct profiles are observed: similar levels of HlgC and HlgB on the left side and higher levels of HlgB compared to HlgC on the right side. For three strains, HlgC and HlgB were below the detection level.

## Results

### High variation in HlgCB production among clinical strains: two distinct profiles

To validate the previous semi-quantitative proteomic analysis of exotoxins (16), the two virulence factors HlgC and HlgB were quantified using isotope-dilution targeted proteomics in a subset of 39 clinical isolates of the previous cohort. The USA300 SF8300 (USA300) and HT20020209-LUG1799 (ST80) strains were used as control strains because their exoprotein profile and virulence in various animal models are well-defined (18,19). Absolute quantification showed considerable HlgC and HlgB variations between isolates with two distinct profiles. In four strains including USA300, the amount of HlgB protein was 6 to 14 times higher than that of HlgC which was very low. In most of the other strains, including ST80 and the PVL-negative PEN strain isolated from a deceased patient (4), similar amounts of HlgB and HlgC were observed, and in the majority of cases the total amount of protein was higher than in the first group (Fig. 1B).

### High levels of HlgCB impacts severity in a rabbit model of pneumonia

To experimentally confirm that high levels of HlgCB increases the virulence of *S. aureus* and the mortality rate of CAP, the PEN strain was tested in a rabbit model of pneumonia. Its virulence was compared to USA300 and ST80, both highly lethal PVL-positive MRSA strains (18,19). All infected rabbits died within 48 h (Fig. 2A) with high lung bacterial densities and high lung weight to body weight ratio (LW/BW) (Fig. 2B and 2C). To determine whether the virulence of PEN was caused by its high production of HlgCB, a premature stop-codon was inserted in the *hlg*C gene (PEN<Q63X) to prevent the production of HlgC and thus HlgCB. Proteomics indicated that HlgB was still produced in PEN<Q63X, albeit to a lower level than in the PEN WT strain (Supplementary Table S1). Rabbits infected with PEN<Q63X showed significantly greater survival (Fig. 2A) and less severe bacterial density and LW/BW ratio (Fig. 2B and 2C) than rabbits infected with PEN WT. In addition, no inflammatory response was observed in rabbits infected by PEN<Q63X, whereas similar levels of IL-1β and IL-8 were quantified in rabbits infected by USA300, ST80, and PEN WT strains (Fig. 2D). These data strongly demonstrate that, *in vivo*, high levels of HlgCB in PVL-negative strains correlate with severe outcome and mortality.

**Figure 2.**
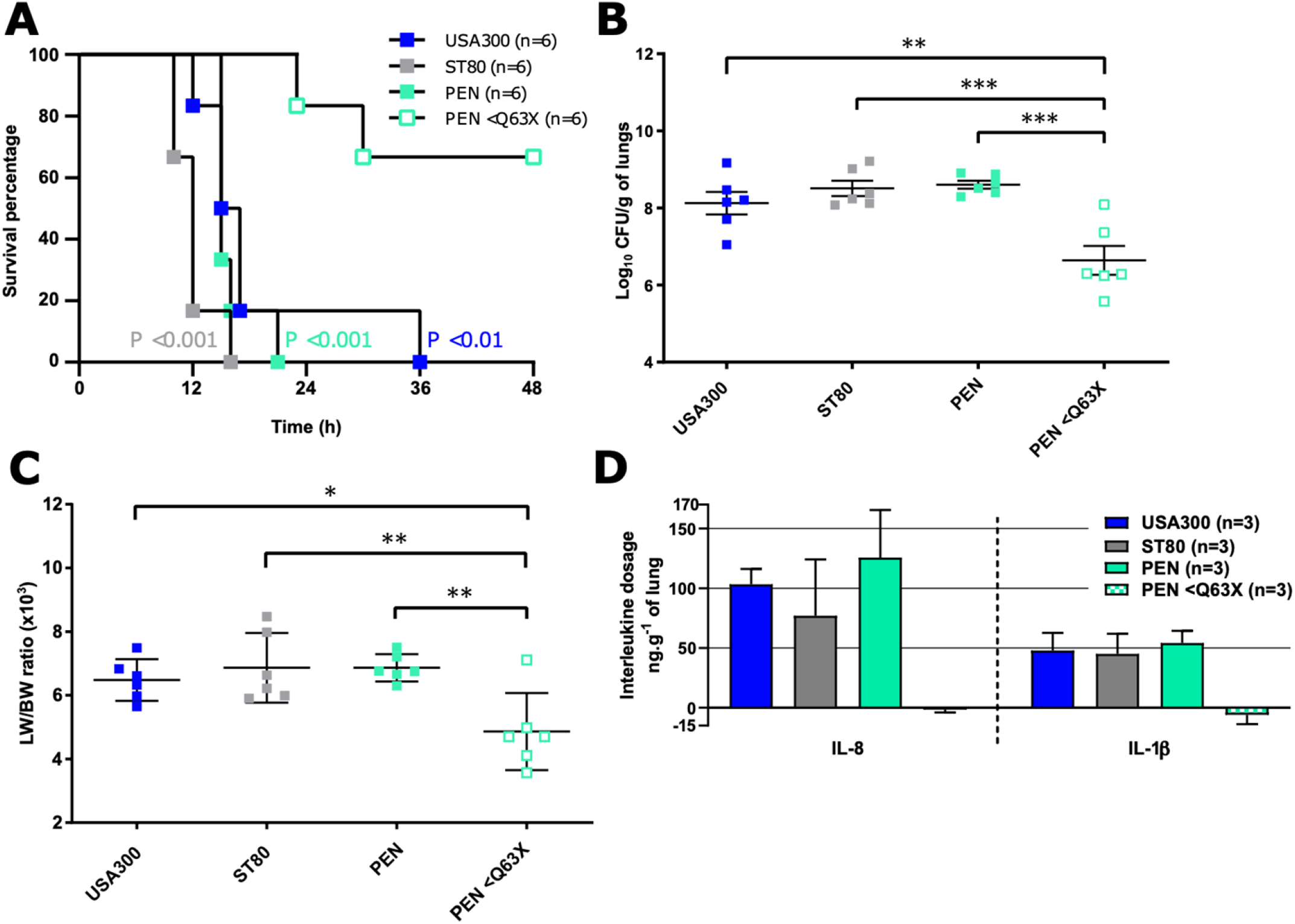
HlgCB contributes to *S. aureus* virulence during pneumonia. (A) Kaplan-Meier survival curves for rabbits infected by endotracheal instillation of 9.5 to 9.6 log10 colony-forming units (CFU) /mL of USA300, ST80, PEN, or PEN with a premature stop-codon in hlgC (PEN <Q63X) strains, to induce necrotizing pneumonia. Log-rank (Mantel-Cox) test was used to compare mortality of PEN HlgC<Q63X to that of USA300 (dark blue), ST80 (grey), and PEN (green). (B and C) Bacterial densities in the lung on a log10 scale (B) and lung weight to body weight ratio (LW/BW) × 10^3^ (C) at the time of death were compared by ANOVA analysis. *P < 0.05; **P < 0.01; ***P < 0.001; non-significant comparisons are not shown. (D) ELISA quantification of IL-8 and IL-1β in lung shred samples collected from rabbits infected with USA300, ST80, PEN, and PEN HlgC<Q63X strains. Quantifications were reported relative to total lung mass.

### Analysis of *hlgCB* operon reveals a processed *hlg*B transcript

To correlate the proteomic profiles of USA300, ST80, and PEN strains with RNA levels, Northern blot analysis was performed on the three strains using specific probes complementary to *hlg*A, *hlg*C, and *hlg*B mRNAs. The *hlg*C and *hlg*B probes unveiled the expected bi-cistronic *hlg*CB transcript (Fig. 3A). A significant variation in *hlg*CB expression was observed, with PEN producing more *hlg*CB mRNA than USA300 and ST80. As expected, no correlation between *hlg*CB and *hlg*A mRNA expression was found. Indeed, both USA300 and PEN yielded more *hlg*A mRNA than ST80, although ST80 produced more *hlg*CB mRNA than USA300. These data suggest that the bi-cistronic *hlg*CB mRNA is transcribed under its own promoter. In addition, the *hlg*B-specific probe revealed the presence of a smaller product, the size of which corresponded to *hlg*B alone. Moreover, *hlg*B transcript level was much higher than that of *hlg*CB in USA300, while the yields of the two transcripts were roughly equivalent in ST80 and PEN strains.

**Figure 3.**
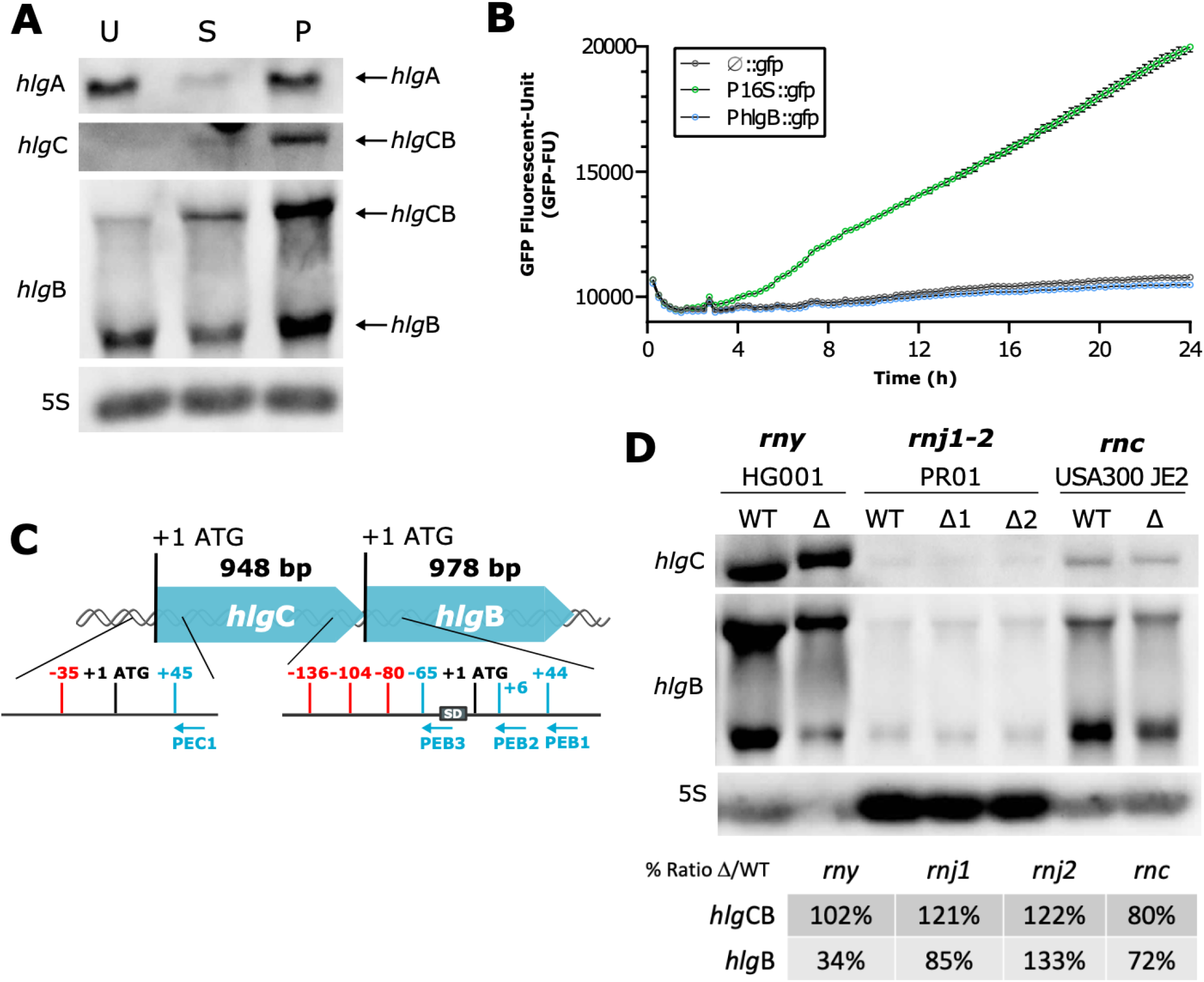
Different expression levels of *hlg*A and *hlg*CB mRNA in three clinicals strains: an additional *hlg*B transcript, produced from the processing of *hlg*CB mRNA. (A) Northern blot analysis of the gamma-hemolysin expression in USA300 (U), ST80 (S), and PEN (P) strains. Total RNA were extracted from 8 h CCY cultures. Probes targeting *hlg*A, *hlg*C, and *hlg*B mRNA were used and are indicated on the left side. The *hlg*B probe detected two transcripts: *hlg*CB mRNA and *hlg*B mRNA, as mentioned on the right side. 5S rRNA (5S) was used as a loading control. (B) The existence of an *hlg*B-specific promoter within the *hlg*C gene was tested using transcriptional fusions in the USA300 strain. The 926bp region upstream *hlg*B and including *hlg*B ATG was cloned into the pACL-1484 plasmid in front of the *gfp* gene (P*hlg*B::*gfp*). The 16S rRNA promoter was used as positive control (P16S::*gfp*) and the plasmid with no insert as a negative control (ø::*gfp*). Promoter activity was measured by GFP signal quantification (GFP Fluorescence Unit –GFP-FU) overtime, during a 24 h kinetic, with a 15 min interval time in CCY medium containing chloramphenicol. Mean with standard-error from biological triplicates are represented in grey for pACL-ø, in green for pACL-P16S, and in blue for pACL-*hlg*B. (C) Schematic summary of primer extension results. Primer extension was performed on total RNA from USA300 and ST80 strains using 5’-radiolabeled primers complementary to *hlg*C (PEC1) and *hlg*B (PEB1, PEB2, and PEB3) sequences. All identified +1 are in red, primers are in blue, and *hlg*B Shine-Dalgarno (SD) is represented with a grey box. (D) Northern blot analysis of *hlg*CB and *hlg*B transcripts in wild type (WT) and mutant (Δ) strains of the main *S. aureus* RNases. Total RNAs were prepared after 8 h in CCY culture of HG001 WT strain and Δ*rny* (RNase Y) mutant, PR01 WT strain and Δ*rnj*1 and Δ*rnj*2 (RNases J1 and J2) mutants, and USA300 JE2 WT strain and Δ*rnc* (RNase III) mutant, and run on 1% agarose gels. After transfer onto a nitrocellulose membrane, *hlg*C and *hlg*B mRNAs were revealed after hybridization of the membrane with specific probes. 5S rRNA (5S) was used as a loading control and was run on a separate gel in parallel to the experiment. Mutant (Δ)/WT ratios are given as percentages of the signal quantified and normalized on 5S signal.

We then investigated whether the *hlg*B transcript was under the control of a specific promoter upstream *hlg*B. The 932 bp upstream the *hlg*B start codon of USA300 and ST80 strains (the sequence of PEN being strictly identical to USA300) were cloned into the pALC1484 plasmid in front of the *gfp* gene. GFP signal quantification failed to detect any promoter activity, as demonstrated by the absence of signal for both the *hlg*B-upstream region tested (pALC-P*hlg*B) and the empty vector (pALC- ø), unlike the positive control (pACL-P16S) containing the 16S-encoding gene promoter (Fig. 3B and Supplementary Fig. S1). The transcriptional starts of *hlg*B and *hlg*CB transcripts were then characterized by primer extension in ST80 and USA300 (Supplementary Fig. S2B). The results showed three possible start sites located at −80, −104, and −136 upstream the *hlg*B start codon whereas only one stop positioned at −35 upstream *hlg*C was found (Fig. 3C and Supplementary Fig. S2). Since *bona fide* mRNA can be distinguished from processed RNA by the presence of a 5’-triphosphate, the Terminator™ 5’-phosphate-dependent exonuclease, which preferentially digests RNA carrying a 5’-monophosphate, was then used. Northern blot experiments showed full degradation of the short *hlg*B mRNA for the three strains, reinforcing the involvement of a processing event. Surprisingly, the *hlg*CB mRNA was also sensitive to the exonuclease treatment while the control *bona fide* sRNA RsaI was resistant (Supplementary Fig. S3). Finally, we explored the impact of the major RNases on the generation of *hlg*B mRNA using single mutant strains deleted of the genes encoding either RNase Y, RNase J1, RNase J2, or RNase III (Fig. 3D). Deletion of RNase Y caused a reproducible reduction in *hlg*B RNA level (66%) as did the Δ*rnc* mutant strain to a lesser extent (28%). In the PR01 genetic background, even though the *hlg*CB and *hlg*B levels were rather low, the *rnj*1 or *rnj*2 deletion had no noticeable effect. Altogether, these data suggest that the *hlg*B transcript results from the processing of the *hlg*CB transcript, which probably involves several RNases including RNase Y.

### Promoter activity and mRNA stability impact *hlg*CB expression

We then assessed the strength of the *hlg*CB promoter and the mRNA stability of the *hlg*CB transcript in the three prototypic strains. A fragment of 434 bp upstream the *hlg*C start codon (P*hlg*C) from USA300, ST80, and PEN strains (Supplementary Fig. S4), was cloned in front of the *gfp* gene in plasmids that were introduced into the three genetic backgrounds. The P*hlg*C of PEN was significantly stronger than the P*hlg*C of the two other strains regardless of the genetic background in which the plasmid was expressed. The P*hlg*C of USA300 and ST80 had lower but similar strengths in PEN and ST80 backgrounds while the P*hlg*C of USA300 was slightly stronger than the P*hlg*C of ST80 in the USA300 background (Fig. 4A).

**Figure 4.**
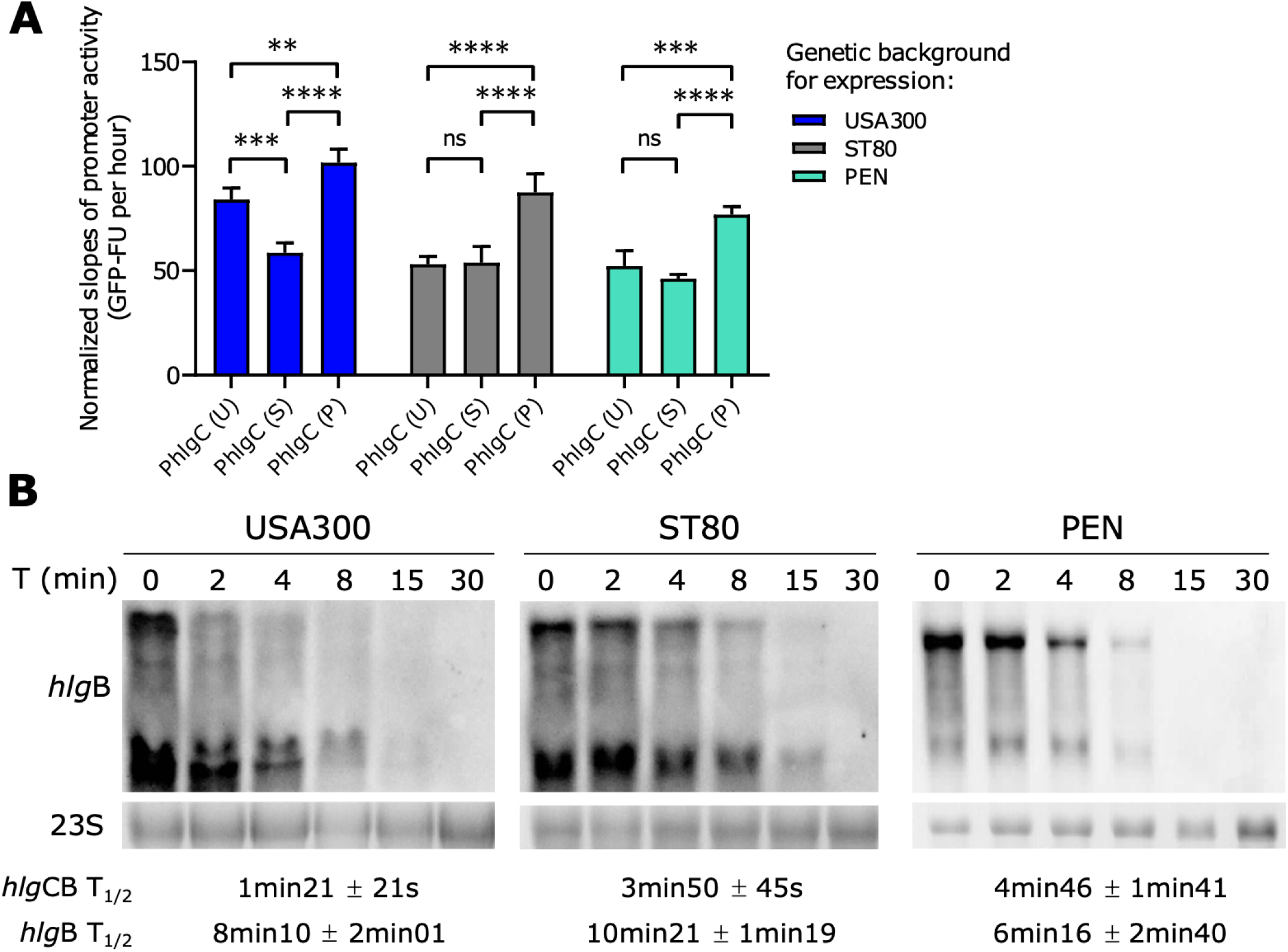
Both promoter activity and mRNA stability/degradation impact *hlg*CB expression. (A) The *hlg*C promoter (P*hlg*C) activity was tested according to its original genetic background and when expressed in autologous and heterologous genetic backgrounds. The 434 bp region upstream and including *hlg*C ATG from USA300 (P*hlg*C (U)), ST80 (P*hlg*C (S)) and PEN (P*hlg*C (P)) strains were cloned into the pACL-1484 plasmid in front of the *gfp* gene. Plasmids were transduced into the three strains: USA300, ST80, and PEN. Promoter activity was measured through GFP signal quantification overtime (GFP Fluorescence unit – GFP-FU), during a 24 h kinetic in CCY. The promoter activity slopes were estimated between the 4 h and 18 h time points, from the mean of biological triplicates, and normalized with negative control. Tuckey’s multiple comparisons test was performed, ns – P_adj_ > 0.05; ** – *P*_adj_ < 0.01; *** – *P*_adj_ < 0.001; and **** – *P*_adj_ < < 0.0001 (B) Determination of *hlg*CB and *hlg*B mRNA stability upon rifampicin exposure. Culture of USA300, ST80, and PEN strains in CCY were treated with rifampicin at 5 h of growth. Total RNA was extracted prior rifampicin addition (T = 0 min) and from 2 min to 30 min after rifampicin addition (T (min)). *hlg*CB and *hlg*B mRNAs were probed with the *hlg*B probe. Ethidium bromide-stained 23S rRNA (23S) was used as a loading control from the same gels. The calculated half-lives (T_1/2_) of *hlg*CB and *hlg*B mRNA relative to 23S are presented below the gel pictures and represent the mean of biological triplicates.

The *hlg*CB and *hlg*B mRNA stabilities were then determined for the three strains using rifampicin assay. Quantification showed that *hlg*B transcript is more stable than *hlg*CB in all strains with noticeable variations between the strains. In USA300, the *hlg*CB mRNA is far less stable than *hlg*B (half-lives: 1 min 21 compared to 8 min 10, respectively) and is also less stable than *hlg*CB from ST80 and PEN strains (3 min 50 and 4 min 46, respectively). The difference between *hlg*CB and *hlg*B is smaller for ST80 (3 min 50 and 10 min 21, respectively) and PEN strains (4 min 46 and 6 min 16, respectively; Fig. 4B).

Altogether, promoter activity and *hlg*CB mRNA stability are characteristic of each genetic background. PEN produces more *hlg*CB mRNA through its strong promoter and has a similar mRNA stability than *hlg*CB in ST80, whereas USA300 presents a weaker promoter than PEN and its *hlg*CB mRNA has a lower stability than the corresponding mRNAs in ST80 and PEN strains.

### An SNP drastically changes *hlg*CB translatability independently of the genetic background

We then questioned whether the polymorphisms in nucleotide sequences of *hlg*CB mRNA may impact ribosome binding and translation efficiency. USA300 and PEN strains only have a 1 nt difference in their 5’UTR sequence at position −13, close to the ribosome binding site. Of note, ST80 has numerous SNPs in the *hlg*C coding region compared to USA300 and PEN but shares the same polymorphism as PEN in the 5’UTR (Fig. 5A).

**Figure 5.**
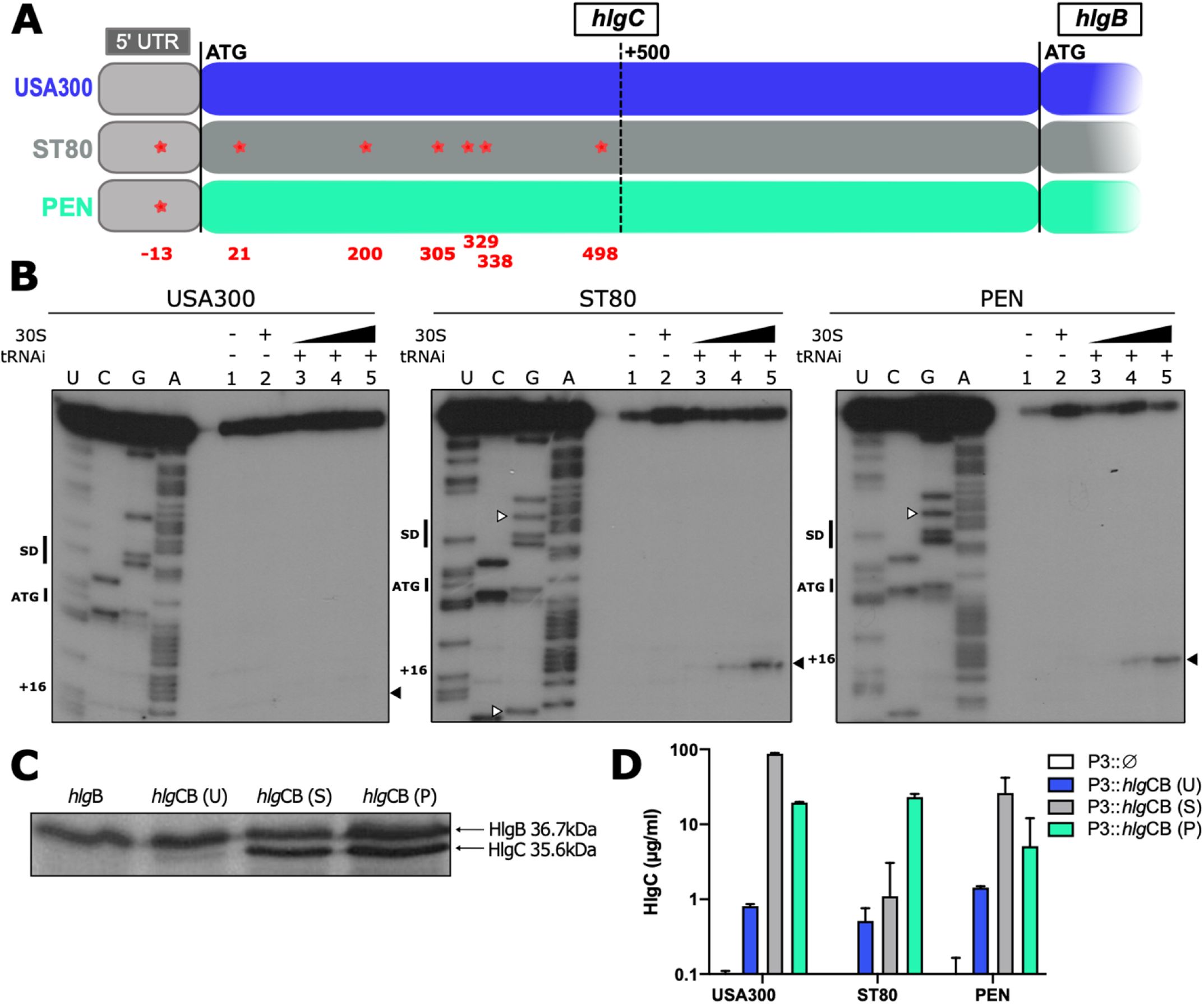
A single SNP drastically changes *hlg*CB translatability, independently of the genetic background. (A) Schematic representation of *hlg*C sequence up to *hlg*B with its 5’-UTR (light grey) from USA300 (blue), ST80 (dark grey), and PEN (green) strains. SNPs between USA300 and ST80 or PEN strains are represented by red stars and nt distance from ATG is mentioned in red under the gene representation. PEN and ST80 strains share the same SNP in the 5’-UTR. (B) Toe-print assays showing binding of *S. aureus* ribosomes on *hlg*CB transcript in the 5’-UTR of *hlg*C mRNA, according to the sequence (USA300, ST80, or PEN). Lane 1: incubation control of mRNA alone; Lane 2: incubation control of mRNA with 30S subunits; Lane 3-5: formation of the ribosomal ternary complex containing mRNA (1 nM), the initiator tRNA^fMet^ (tRNAi, 1 μM), and increasing concentrations of 30S: 0.5 μM (Lane 3), 1 μM (Lane 4), and 2 μM (Lane 5). Lanes U, C, G, and A: sequencing ladders. The Shine and Dalgarno (SD) sequence, the translation start site (ATG), SNPs compared to USA300 sequence (white arrow), and the toe-printing signal (+16) are indicated. (C) *In vitro* translation of *hlg*B and *hlg*CB mRNA using pUC-T7 constructions with *hlg*B or *hlg*CB DNA sequences from USA300 (U), ST80 (S), and PEN (P) strains. *S. aureus* 70S ribosomes were used and the reaction was performed during 4 h with ^35^S-Methionine. (D) Quantification by ELISA (n=3) of HlgC in μg/mL from 8 h CCY culture supernatants. *hlg*CB from USA300 (P3::*hlg*CB (U)), ST80 (P3::*hlg*CB (S)),and PEN (P3::*hlg*CB (P)) strains were cloned into the pCN38 plasmid under the P3 promoter, and overexpressed in USA300Δ*hlg*, ST80 Δ*hlg*, and PEN Δ*hlg* strains. The plasmid pCN38 without *hlg*CB (P3::Ø) was used as negative control.

Formation of a simplified initiation complex was analyzed using toe-printing assays. The experiments were performed using *S. aureus* 30S subunits, the initiator tRNA^Met^, and *in vitro* transcribed *hlg*CB and *hlg*B mRNAs of the three strains. Using an oligonucleotide specific to *hlg*C, a clear toe-print was observed at position +16 from the initiation codon of ST80 and PEN *hlg*C transcripts, but no signal was detected for USA300 mRNA (Fig. 5B). A similar experiment was performed using an *hlg*B specific oligonucleotide to investigate the accessibility of *hlg*B ribosome binding site characterized by a strong Shine-Dalgarno sequence. A clear toe-print at position +16 was observed for all *hlg*CB mRNAs (Supplementary Fig. S5A) and for the *hlg*B transcript (Supplementary Fig. S5B). These data show that the ribosome binding site of *hlg*B mRNA is accessible to promote the formation of a translation initiation complex whatever the mRNA context.

*In vitro* translation assays were then performed using *S. aureus* 70S ribosomes and *hlg*CB mRNA of the three strains or *hlg*B mRNA of ST80, as there is no polymorphism in *hlg*B. The results clearly showed that the *hlg*B and the *hlg*CB transcripts can produce HlgB protein. For HlgC, the amount of protein synthesized was much lower from USA300 *hlg*CB mRNA compared to PEN and ST80 transcripts (Fig. 5C). Hence, HlgB and HlgC levels appeared almost equimolar among ST80 and PEN *hlg*CB mRNAs, but not for USA300 *hlg*CB mRNA. These results confirm the impact of the 5’UTR SNP on translatability and show that *hlg*B can be translated independently from both *hlg*CB and *hlg*B mRNAs *in vitro*. Because the translation of *hlg*C is impaired in USA300, the synthesis of HlgB (Fig. 5C) resulted most probably from an internal entry of the ribosome at the *hlg*B initiation site or from the processed form of *hlg*B as shown by the toe-printing assays (Supplementary Fig. S5).

In order to validate the importance of the 5’UTR SNP on translation efficiency *in vivo*, the *hlg*CB coding sequence from each strain was placed under the control of the P3 promoter and introduced into the three different genetic backgrounds deleted for *hlgABC* genes. As expected, the USA300 sequence, irrespective of the genetic background, produced low levels of HlgC (Fig. 5D). Conversely, both the ST80 and PEN sequences allowed efficient HlgC synthesis. The ST80 sequence under the P3 promoter led to a greater amount of HlgC than PEN sequence, most probably due to its slightly higher RNA stability (Fig. 5D). Overall, the presence of a single SNP in the 5’UTR of *hlg*C has a strong consequence on ribosome recruitment and translation efficiency.

## Discussion

In this study, we first demonstrated that the PVL-negative strain PEN was as virulent as the PVL-positive strains USA300 and ST80, and that this virulence was mainly driven by HlgCB production. When highly produced, HlgCB leads to rapid mortality, strong production of cytokines in the lung, and high bacterial load in the rabbit model of pneumonia, similarly to what is observed with PVL (18,19). Consistently, although HlgCB targets the same receptors as LukS-PV, it interacts with different motifs, leading to better cell permeabilization (12). Since all *S. aureus* strains possess the *hlg* locus (20,21) but not all cause severe CAP (4), the present study strengthens the hypothesis that expression matters (16).

To decipher the mechanisms leading to a variation in HlgCB expression and production among clinical strains, we investigated various levels of regulation including promoter activity, mRNA stability, and translatability. Differences in promoter activity were expected due to the polymorphisms between PEN and ST80 (2 SNPs) or USA300 strains (2 SNPs and 1 insertion) (Supplementary Fig. S4). A variation in expression due to differences in promoter sequences has already been reported for other toxins such as the α-toxin (22). Surprisingly, the promoter activity of P*hlg*C from USA300 was weaker when expressed in the two other strain backgrounds, suggesting an indirect effect caused by the modulation of an unknown transcriptional regulator. In addition, the stability and translatability of mRNA rely on its sequence and resulting structure; in USA300, the weak efficiency of *hlg*C translation probably resulted in an mRNA more vulnerable to degradation (23), leading to almost no production of HlgC. Strikingly, the difference in translation efficiency between USA300, PEN, and ST80 strains is related to a single SNP in the 5’UTR of the *hlg*CB operon. Other studies have shown that the introduction of mutations in the 5’UTR of mRNAs in *E. coli* and *B. subtilis* can induce changes in the mRNA structure with consequences on ribosome recruitment at initiation sites and on translation efficiencies (24–28). Secondary structure prediction of *hlg*C 5’UTR revealed a hairpin structure that partially sequesters the weak SD sequence (Supplementary Fig. S6). The G-13-U mutation in USA300 could give rise to an additional base pair that would reinforce the stability of the hairpin structure. Interestingly, the three strains that have an HlgCB profile similar to USA300, carry either the same SNP at position −13 (FAL003 and MAS222) or an SNP at position −7 (BES288). For BES strain, the SNP at position −7 (G to U) reduces the strength of the SD, and an additional base pair can be formed (Supplementary Fig. S6). It is thus tempting to propose that these two SNPs (−13 and −7) would decrease helix breathing to hinder ribosome binding and/or decrease SD efficiency. In addition, ST80 contains 6 additional SNPs in the coding sequence that do not alter the sequence of HlgC, and thus are considered to be silent. However, a recent study performed in yeast revealed that both synonymous and non-synonymous mutations in coding sequences resulted in a significant reduction in fitness and that the mutations frequently alter mRNA levels (29). Therefore, we do not exclude that the SNPs in the coding sequence might have consequences on the translation speed and/or mRNA stability in ST80. Considering all the data obtained herein, we propose the following scenario to explain the variation in HlgCB levels among clinical strains (Fig. 6). The PEN strain has a strong promoter leading to high amount of translatable and stable *hlg*CB mRNA, allowing a high production of HlgCB. ST80, despite a weaker *hlg*C promoter than PEN, shows better mRNA stability and translatability than USA300 and allows a large production of HlgCB. Both ST80 and PEN strains thus produce high levels of HlgC and HlgB. In contrast, USA300 has an intermediate *hlg*C promoter activity, a SNP in the 5’-UTR of *hlg*C mRNA that prevents *hlg*C translation, and a more unstable *hlg*C mRNA, resulting in very low levels of HlgC (Fig. 6). Altogether, these three levels of *hlg*C regulation contribute to the protein levels measured by absolute quantification in the three strains. Interestingly, although several SNPs between the three strains in and upstream the *hlg*C sequence are observed and responsible for the variation in HlgC production, no SNP was identified in *hlg*B and its protein level was less affected among clinical strain. This might be due to a dual evolutionary constraint, as HlgB also forms heterodimers with HlgA. The high variability in core genome gene expression raises the question of the evolutionary pressure that led to the selection of this expression diversity, which may represent a fine and specific adaptation to the context, such as interactions with the host in which each strain develops. It is very likely that other determinants modulate the overall level of HlgCB production in clinical strains, such as transcriptional or post-transcriptional regulators. As illustrated in this study and elsewhere, epistasis effects as well as the non-coding and coding sequences, all play roles in the expression of virulence factors (30–32). The continuous increase in GWAS (31,33,34) performance for the analysis of large sets of strains may allow, in the future, the identification of these other contributors.

**Figure 6.**
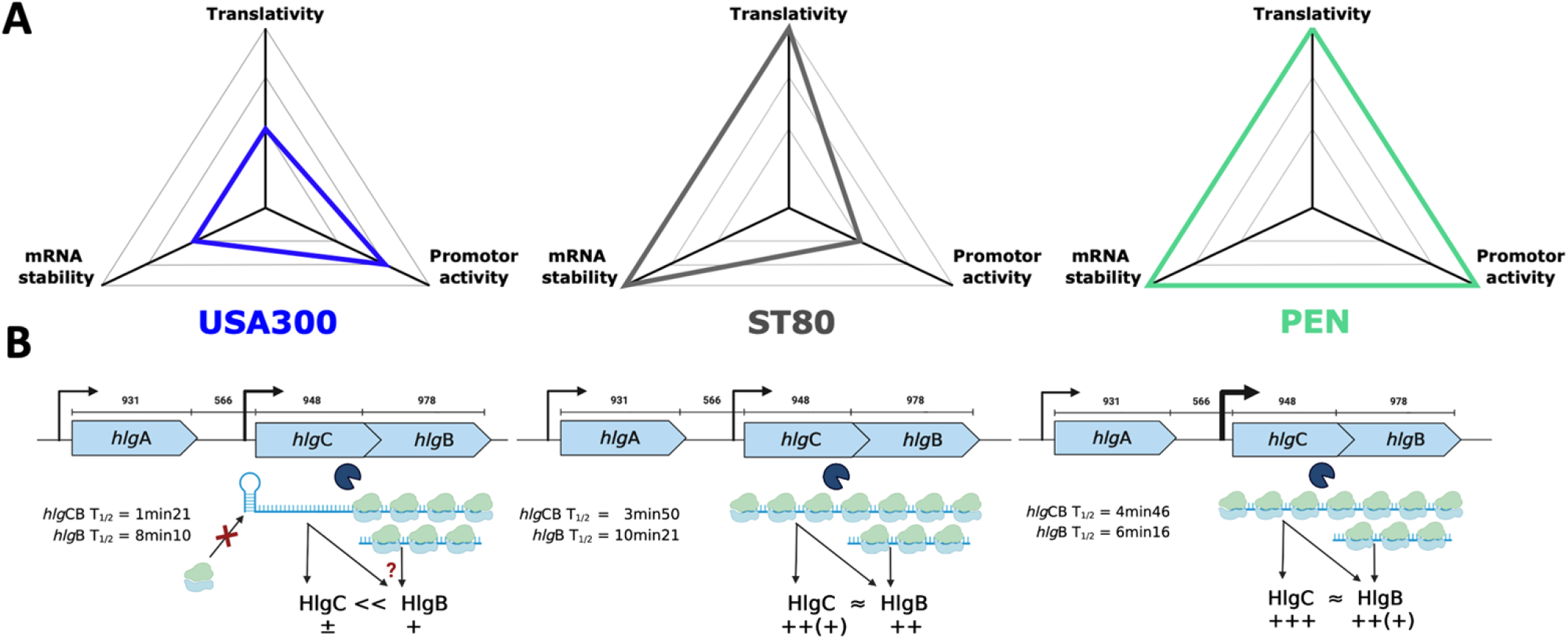
Regulatory mechanisms impacting gamma-hemolysin production. (A) Kiviat diagram representing the different mechanisms explored in this study, and their importance regarding HlgC production in USA300 (blue), ST80 (grey), and PEN (green) strains, with an arbitrary scale unit. The larger the area is, the greater the amount of HlgC will be. For HlgB, the maturated *hlg*B mRNA modifies this correlation between the three mechanisms and the production, especially for USA300 strain. (B) Model for the different levels of regulation of HlgC and HlgB production. In USA300 strain, HlgC production is impaired due to an SNP impeding ribosome recruitment at the *hlg*C initiation site. Translational coupling might thus be abolished and HlgB synthesis might come from the internal entry of the ribosome at the initiation site of *hlg*B on *hlg*CB mRNA or from the processed transcript.

The discovery of a new *hlg*B transcript in this study adds to the overall complexity. Since no promoter activity was detected upstream *hlg*B and the three different 5’ ends carry a 5’-monophosphate, this new transcript is not a *bona fide* mRNA and the maturation process likely involves several RNases. Previous RNA sequencing performed on different strains showed the presence of an antisense RNA against *hlg*C and more *hlg*B than *hlg*C transcript, (35) supporting the existence of a processed form of *hlg*B. Whether the antisense RNA is involved in the maturation process of *hlg*B remains to be determined. The processed *hlg*B mRNA appears to be translated *in vitro* and, in the case of USA300 where *hlg*C translation is impeded leading to an unstable mRNA, the synthesis of HlgB might originate from the *hlg*B mRNA. As the HlgB component is used for both HlgCB and HlgAB toxins (36), the *hlg*B transcripts could be translated under specific conditions to allow the production of HlgB independently of HlgC, in order to form the HlgAB toxin. Unlike *hlg*C and *hlgB, hlg*A is encoded upstream *hlg*CB operon and possesses its own promoter (13,37). Indeed, there was no correlation herein between *hlg*A and *hlg*CB mRNAs levels, an observation supported by previous studies in which *hlg*A and *hlg*CB expression differed from each other (38). On one hand, *hlg*CB is up-regulated by the *agr* system (38,39), SarA (38,39), and SaeRS regulation system (40–44) but down-regulated by *rot* (45). On the other hand, *hlg*A is mostly regulated by the SaeRS system(42,43,46,47), but not by *agr* nor SarA (38,46). The dissociated regulation of *hlg*A and *hlg*B remains unexpected since HlgA requires HlgB to induce pore-formation (48). One hypothesis is that the *hlg*B transcript might be appropriate in specific conditions to optimize the formation of the HlgAB toxin. To further complexify the matter, non-equimolar productions of HlgA and HlgB may be relevant in certain conditions, as HlgA alone antagonizes its receptors (CXCR1, CXCR2 and CCR2) preventing neutrophil activation (11).

This work illustrates the complexity of virulence factor expression in clinical strains, where a given gene can produce a protein at highly variable levels not only due to the corresponding promoter activity. Furthermore, proteins from an operon may not be expressed equimolarly, and proteins belonging to different operons but who are supposed to form an equimolar assembly may not be produced at equivalent levels. It is very likely that such phenomena will be discovered in other bacterial species as they probably participate in the evolutionary plasticity of bacteria.

## Material and Methods

### Bacterial strains and growth conditions

All strains and plasmids used in this study are listed in Supplementary Table S2. *S. aureus* strains were routinely cultured overnight in Tryptic Soy Broth (TSB, DIFCO), then diluted at OD_600_=0.05 in CCY (3% wt/vol yeast extract, 2% bacto-casamino acids, 2.3% sodium pyruvate, 0.63% Na_2_HPO_4_, and 0.041% KH_2_PO_4_) and grown for 5 h (Rifampicin test) or 8 h at 37°C with shaking. When required, chloramphenicol was added at the concentration of 10 μg/mL.

For animal experiments, strains were inoculated into 5 mL of CCY medium and incubated overnight at 37°C with shaking. Then, 500 μL of this culture were inoculated in CCY and agitated for 4 h at 37°C to reach the exponential growth phase.

### Construction of mutant strains

The *S. aureus hlgC* stop codon mutant was created in the clinical PEN strain using the pnCasSA-BEC genome editing system as previously described (49) (Supplementary M&M). The entire *hlg*ACB locus was also deleted in USA300, ST80, and PEN strains by double recombination process as previously described (50) (Supplementary M&M).

### Absolute quantification of HlgC and HlgB in clinical strains using targeted proteomics

*S. aureus* clinical strains were cultured 8 h in CCY medium and centrifuged at 10 000 rpm for 5 min at 4°C to separate bacterial pellets. For each strain, 50 μl of supernatant (50 μg of proteins) were reduced, alkylated, and digested using LysC/trypsin mix (Promega) (51). To quantify HlgC and HlgB, two signature peptides were selected and purchased in an isotopically-labeled version (AQUA QuantPro, Thermo Fisher Scientific) to serve as quantification standards: YVSLINYLP[^13^C_6_, ^15^N_2_]K (HlgC) and SNFNPEFLSVLSH[^13^C_6_, ^15^N_4_]R (HlgB). These labeled peptides were added to the digested supernatant at a final concentration of 2.1 pmoles/mL. Then, the digest was desalted on a C18 ZipTip (MacroSpin Harvard) before drying by vacuum centrifugation and storage at −20°C. Dried peptide digest was solubilized in 40 μL of 2% acetonitrile and 0.1% formic acid, and 6 μL of this solution was analyzed via targeted proteomics. Targeted proteomic analyses were performed on a 6500 QTrap mass spectrometer (AB Sciex) operating in the selected reaction mode (SRM). Liquid chromatography (LC) separation was performed on an Ultimate 3000 system (Thermo Scientific). The parameters for LC separation and scheduled SRM acquisitions are presented in the Supplementary M&M. LC-SRM data analysis was performed using Skyline software. All transitions were individually inspected and excluded if they were deemed unsuitable for quantification (low signal-to-noise ratio, obvious interference). Unlabeled/labeled peak area ratios were calculated for each SRM transition and these ratios were used to determine the corresponding mean peptide ratio. HlgC and HlgB concentrations in *S. aureus* supernatants were calculated based on the mean ratio of each signature peptide.

### Rabbit model of pneumonia

Immunocompetent New Zealand White male rabbits (body weight 2.8 to 3.2 kilograms) were challenged at Vivexia (Dijon, France) by an intrabronchial instillation of 0.5 ml of *S*. *aureus* strains diluted at 9.49 to 9.61 log_10_ CFU/mL in sterile saline solution (Otec^®^, NaCl 0,9%) to induce pneumonia. The inoculum was gently flushed through the tracheal catheter under laryngoscopic control. The catheter was immediately removed, the animals were rapidly extubated, and allowed to go back to their cage. Post-infection rabbits’ survival rate was then recorded 2 to 3 times a day. If they had not spontaneously died, animals were euthanized 48 h after the challenge. Lungs were collected for cytokine and bacterial load estimation: each lung (right and left) was weighed and homogenized in 10 mL of sterile saline solution. Bacterial densities were counted in crude lung homogenates by plating 10-fold dilutions on Chapman agar and incubating the plates for 24 h at 37°C. Bacterial concentrations were expressed as log_10_ CFU/g. All procedures in the protocol were approved by the local ethics committee for animal experiments and by the *Ministère de l’Enseignement Supérieur et de la Recherche* (APAFIS#22710-201911071356264 v1).

### Cytokine quantification

IL-8 and IL-1β cytokines were quantified by ELISA using the Rabbit IL-1 beta ELISA Kit and Rabbit IL-8 ELISA Kit (Invitrogen). Aliquots of 2 mL of lung homogenate shred in saline solution were centrifuged during 30 min at 10 000 rpm and 4°C, before appropriate dilution in Assay Diluent solution. ELISA were then performed by strictly following the supplier’s protocols. The final concentration was reported on the total mass of the lung. Three biological replicates were used for each strain tested.

### Northern blot analysis

Total RNA was prepared from 10 mL of 8 h culture following the procedure described for the FastRNA pro blue kit (MP Biomedicals) with the Fastprep apparatus (MP Biomedicals). Electrophoresis of total RNA (10 to 30 μg) was performed on 1.5% agarose gel containing 25 mM guanidium thiocyanate. After migration, RNAs were transferred by capillarity on nitrocellulose membranes (GE Healthcare Life Sciences). Hybridization with specific digoxygenin (DIG)-labelled probe complementary to *hlg*A, *hlg*C, *hlg*B, RsaI, and 5S rRNA sequences (Supplementary Table S3) followed by luminescent detection were carried out as previously described (52).

### Primer extension assays

To determine the transcriptional start sites of *hlg*CB and *hlg*B mRNAs, primer extension assays were performed as previously described (53). Briefly, 30 μg of total RNAs from USA300 and ST80 strains were reversed transcribed with AMV reverse transcriptase (NEB) using 5’ radiolabelled primers (for *hlg*C (PEC1) and for *hlg*B (PEB1, PEB2, and PEB3); Supplementary Table S3). The sequencing ladders were obtained from *hlg*CB or *hlg*B PCR products (amplified with primers listed in Supplementary Table S2) and the Vent (exo-) DNA polymerase (NEB Biolabs). All samples were fractionated on a 10% polyacrylamide −8 M urea gel in 1x TBE. After migration, gel was exposed to an autoradiography film.

### Promoter activity assays

The regions upstream *hlg*C (434 bp) and *hlg*B (929 bp) and including their start codons, were amplified from USA300, PEN, and ST80 gDNAs for *hlg*C and only ST80 gDNA for *hlg*B (because no sequence differences were observed between the three strains) using oligonucleotides PhlgC-*EcoR*I-F with PhlgC-*Xba*I-R and PhlgB-*Puv*II-F with PhlgB-*Kpn*I-R for *hlg*C and *hlg*B upstream regions, respectively (Supplementary Table S2). Then, amplicons were fused to the *gfppuv* gene by ligation in pALC1484 plasmid (54) using adequate restriction enzymes (Supplementary Table S3). The 16S rRNA promoter was cloned into pACL1484 as a positive control, as previously described (55). All constructions were verified by PCR and sequencing using pALC-F and pACL-R oligonucleotides (Supplementary Table S3). Plasmids were first electroporated in RN4220 strain prior transformation by electroporation in USA300, ST80, and PEN strains. A 96-well plate (Greiner bio-one – 655090) was inoculated with 200 μL per well of OD_600_=0.05 culture in CCY + 10 μg/mL of chloramphenicol and incubated at 37° for 24 h. GFP measurements were performed with Infinite 200Pro (TECAN) apparatus, with an excitation wavelength at 488 nm and an emission wavelength at 535 nm. Both GFP signal and OD_600_ were measured every 15 min for 24 h. Each strain was evaluated in triplicate.

### RNA stability assays

*hlg*CB and *hlg*B mRNA stability were determined by rifampicin assays. Briefly, USA300, ST80, and PEN strains were grown for 5 h in 200 mL of CCY, and treated with 1 mL of rifampicin at 30 mg/mL to inhibit transcription. Total RNA was extracted before (time 0) and after rifampicin addition at 2, 4, 8, 15, and 30 min. Northern blots were performed using 23S rRNA as loading control.

### Toe-printing assays

*hlg*CB and *hlg*B mRNA transcripts were first synthetized from linearized pUC-T7 vectors (56). The *hlg*CB sequences from USA300, ST80, and PEN strains, and *hlg*B from ST80 strain, containing their 5’UTR, were amplified by PCR using pUCT7-*hlg*C- or -*hlg*B-*Stu*I-F with pUCT7-*hlg*B-*BamH*I-R primers (Table S3), containing the T7 promoter. After linearization by *BamH*I and *in vitro* transcription using T7 RNA polymerase, the DNA template was digested with DNase I (Sigma) and the RNAs were purified with phenol-chloroform alcohol isoamyl, precipitated in absolute ethanol and cleaned-up with Micro Bio-Spin Chromato Column (Biorad) (52). Toe-printing assays were performed on *hlg*CB and *hlg*B transcripts. The RNAs were first annealed to appropriate 5’end-radiolabeled oligonucleotide as previously described (52). The ternary ribosomal complex containing purified *S. aureus* 30S ribosome(57) (0.5, 1, and 2 μM), initiator tRNA^fMet^ (1 μM), and RNA transcripts (1 nM) was formed at 37°C. After reverse transcription, the DNA fragments were sized on 8% polyacrylamide-urea gel electrophoresis. Sequencing ladders were run in parallel to identify the toe-print signals.

### *In vitro* translation assays

The translatability of *hlg*CB and *hlg*B mRNAs were assessed using the PURExpress^®^ Δ Ribosome Kit (NEB) supplemented with purified *S. aureus* 70S ribosomes (57). The ribosomes were first reactivated by incubation at 37°C for 10 min. The reactions were performed 4 h at 37°C with various plasmids (200 ng; pUC-T7::*hlg*CB from USA300, ST80, and PEN strains and pUC-T7::*hlg*B from ST80 strain) and *S. aureus* 70S (30 pmoles) in 5 μL of Buffer A, 1.5 μL of factors, 1 μL of RNasine (Promega), and 1 μL of ^35^S-Methionine. The Laemmli Buffer was added to the samples, which were loaded on 15% PAGE-SDS with Page Ruler™ Plus Protein Product (Thermofisher) run in parallel. After migration, the gel was heat-vacuum transferred on a Whatman paper and the data were revealed by autoradiography.

### Over-expression of *hlg*CB mRNA

The sequence from −39 nt upstream *hlg*C ATG to +60 nt after *hlg*B stop codon was amplified with *hlg*CB-*Pst*I-F and *hlg*CB-*BamH*I-R oligos (Table S3) from gDNA of USA300 (*hlg*CB (U)), ST80 (*hlg*CB (S)), and PEN (*hlg*CB (P)) strains. Then, amplicons were cloned into the pCN38-P3 plasmid using adequate restriction enzymes downstream the P3 promoter. All constructions were verified by PCR and sequencing. Plasmids were first electroporated in RN4220 prior transformation by electroporation in USA300Δ*hlg*, ST80Δ*hlg*, and PENΔ*hlg* strains. All strains were cultivated overnight in TSB with chloramphenicol and then subcultured at OD_600_=0.05 in CCY without antibiotic and incubated during 8 h. Supernatants were centrifuged 5 min at 10 000 rpm at 4°C and stored at −80°C.

### HlgC quantification by ELISA

HlgC quantification was performed by sandwich ELISA using a custom-made mouse monoclonal antibody (R&D Biotech) and a custom-made polyclonal rabbit F(ab)’2 biotinylated antibody (R&D Biotech), both raised against HlgC. A 96-well NUNC Maxisorp plate (Thermo Scientific) was coated with anti-HlgC monoclonal antibody at 10 μg/mL overnight at 20°C. After 5 consecutive washes with PBS Tween 0.05% (PBS-T), wells were saturated 1h30 at 20°C with blocking solution of PBS-T, low fat milk (5 g/L) and BSA (1 g/L). Standard dilutions (15–1000 ng/mL) of recombinant HlgC or culture supernatant were denatured 1 h at 95°C, loaded in duplicate, and incubated for 2 h at 37 °C. Plate was washed, and polyclonal rabbit F(ab)’2 biotinylated antibody (1,55 μg/mL) was added and incubated 1h30 at 37°C. After a new washing step, ExtrAvidin-Peroxidase antibody (Sigma) targeting biotin molecule and peroxidase HRP-conjugated was added and incubated 1 h at 20°C. A final wash was performed prior revelation with 75 μL of substrate tetramethylbenzidine (KPL SureBlue™, SeraCare). The reaction was stopped with H_2_SO_4_ at 1 N. The plates were read at 450 nm in a Biorad Model 680 microplate reader.

## Acknowledgements

We thank Sylvère Bastien for bioinformatics support, Xanthe Adams for technical help, Pr. Christiane Wolz, Pr. Peter Redder and Dr. Paul Verhoven for providing strains, Iuilia Macavei and Jérôme Lemoine for proteomics analysis, and Véréna Landel from the Hospices Civils de Lyon (France) for help in manuscript preparation.

This work was funded by the ANR Ribostaph and by the RHU IDBIORIV.

## References for Supplementary data

(58) (59) (60) (61) (62) (63) (64) (65) (66) (67) (68) (69) (70)

## Supporting Data

### Supplementary material & methods

#### Supplementary figures

- Figure S1: *hlg*B does not have its own promoter
- Figure S2: Primer extension assays: *hlg*B mRNA has three different +1
- Figure S3: *hlg*B mRNA is a maturation product of the *hlg*CB transcript
- Figure S4: Pairwise Sequence Alignment: only few SNPs are detected in the *hlg*C upstream region of USA300, ST80, and PEN strains
- Figure S5: *hlg*B mRNA translation initiation does not require *hlg*C translation
- Figure S6: Secondary structure prediction of *hlg*C 5’UTR including 40 nucleotides from the coding sequence

#### Supplementary tables

- Table S1: Relative quantities of HlgC and HlgB in PEN wild-type and mutant strains by semi-quantitative proteomic analysis
- Table S2: Strains and plasmids used in the study
- Table S3: Oligonucleotides used in the study

## References

1. Thammavongsa V, Kim HK, Missiakas D, Schneewind O. Staphylococcal manipulation of host immune responses. Nat Rev Microbiol. 2015 Sep;13(9):529–43.

2. Tam K, Torres VJ. Staphylococcus aureus Secreted Toxins and Extracellular Enzymes. Microbiol Spectr. 2019;7(2).

3. Jean SS, Chang YC, Lin WC, Lee WS, Hsueh PR, Hsu CW. Epidemiology, Treatment, and Prevention of Nosocomial Bacterial Pneumonia. J Clin Med [Internet]. 2020 Jan 19 [cited 2020 Apr 23];9(1). Available from: https://www.ncbi.nlm.nih.gov/pmc/articles/PMC7019939/

4. Gillet Y, Tristan A, Rasigade JP, Saadatian-Elahi M, Bouchiat C, Bes M, et al. Prognostic factors of severe community-acquired staphylococcal pneumonia in France. Eur Respir J. 2021 Nov;58(5):2004445.

5. Vardakas KZ, Matthaiou DK, Falagas ME. Incidence, characteristics and outcomes of patients with severe community acquired-MRSA pneumonia. European Respiratory Journal. 2009 Nov 1;34(5):1148–58.

6. Masters IB, Isles AF, Grimwood K. Necrotizing pneumonia: an emerging problem in children? Pneumonia (Nathan) [Internet]. 2017 Jul 25 [cited 2020 Apr 6];9. Available from: https://www.ncbi.nlm.nih.gov/pmc/articles/PMC5525269/

7. Gillet Y, Issartel B, Vanhems P, Fournet JC, Lina G, Bes M, et al. Association between Staphylococcus aureus strains carrying gene for Panton-Valentine leukocidin and highly lethal necrotising pneumonia in young immunocompetent patients. Lancet. 2002 Mar 2;359(9308):753–9.

8. Spaan AN, Henry T, van Rooijen WJM, Perret M, Badiou C, Aerts PC, et al. The Staphylococcal Toxin Panton-Valentine Leukocidin Targets Human C5a Receptors. Cell Host & Microbe. 2013 May 15;13(5):584–94.

9. Tromp AT, Van Gent M, Abrial P, Martin A, Jansen JP, De Haas CJC, et al. Human CD45 is an F-component-specific receptor for the staphylococcal toxin Panton-Valentine leukocidin. Nat Microbiol. 2018;3(6):708–17.

10. Grumann D, Nübel U, Bröker BM. Staphylococcus aureus toxins – Their functions and genetics. Infection, Genetics and Evolution. 2014 Jan 1;21:583–92.

11. Spaan AN, Vrieling M, Wallet P, Badiou C, Reyes-Robles T, Ohneck EA, et al. The staphylococcal toxins γ-haemolysin AB and CB differentially target phagocytes by employing specific chemokine receptors. Nature Communications. 2014 Nov 11;5:5438.

12. Spaan AN, Schiepers A, Haas CJC de, Hooijdonk DDJJ van, Badiou C, Contamin H, et al. Differential Interaction of the Staphylococcal Toxins Panton–Valentine Leukocidin and γ-Hemolysin CB with Human C5a Receptors. The Journal of Immunology. 2015 Aug 1;195(3):1034–43.

13. Cooney J, Kienle Z, Foster TJ, O’Toole PW. The Gamma-Hemolysin Locus of Staphylococcus aureus Comprises Three Linked Genes, Two of Which Are Identical to the Genes for the F and S Components of Leukocidin. 1993;61:4.

14. Lubkin A, Lee WL, Alonzo F, Wang C, Aligo J, Keller M, et al. Staphylococcus aureus Leukocidins Target Endothelial DARC to Cause Lethality in Mice. Cell Host Microbe. 2019 Mar 13;25(3):463–470.e9.

15. Dajcs JJ, Thibodeaux BA, Girgis DO, O’Callaghan RJ. Corneal Virulence of Staphylococcus aureus in an Experimental Model of Keratitis. DNA and Cell Biology. 2002 May;21(5–6):375–82.

16. Pivard M, Bastien S, Macavei I, Mouton N, Rasigade JP, Couzon F, et al. Staphylococcus aureus virulence factor expression matters: input from targeted proteomics shows Panton-Valentine leucocidin impact on mortality [Internet]. bioRxiv; 2022 [cited 2022 Sep 20]. p. 2022.09.18.508069. Available from: https://www.biorxiv.org/content/10.1101/2022.09.18.508069v1

17. Stulik L, Rouha H, Labrousse D, Visram ZC, Badarau A, Maierhofer B, et al. Preventing lung pathology and mortality in rabbit Staphylococcus aureus pneumonia models with cytotoxin-neutralizing monoclonal IgGs penetrating the epithelial lining fluid. Sci Rep. 2019 Mar 29;9(1):5339.

18. Diep BA, Chan L, Tattevin P, Kajikawa O, Martin TR, Basuino L, et al. Polymorphonuclear leukocytes mediate Staphylococcus aureus Panton-Valentine leukocidin-induced lung inflammation and injury. PNAS. 2010 Mar 23;107(12):5587–92.

19. Diep BA, Le VTM, Badiou C, Le HN, Pinheiro MG, Duong AH, et al. IVIG-mediated protection against necrotizing pneumonia caused by MRSA. Sci Transl Med. 2016 21;8(357):357ra124.

20. McCarthy AJ, Lindsay JA. Staphylococcus aureus innate immune evasion is lineage-specific: a bioinfomatics study. Infect Genet Evol. 2013 Oct;19:7–14.

21. von Eiff C, Friedrich AW, Peters G, Becker K. Prevalence of genes encoding for members of the staphylococcal leukotoxin family among clinical isolates of Staphylococcus aureus. Diagn Microbiol Infect Dis. 2004 Jul;49(3):157–62.

22. Tavares A, Nielsen JB, Boye K, Rohde S, Paulo AC, Westh H, et al. Insights into alpha-hemolysin (Hla) evolution and expression among Staphylococcus aureus clones with hospital and community origin. PLoS One. 2014;9(7):e98634.

23. Dar D, Sorek R. Extensive reshaping of bacterial operons by programmed mRNA decay. PLoS Genet. 2018 Apr 18;14(4):e1007354.

24. de Smit MH, van Duin J. Secondary structure of the ribosome binding site determines translational efficiency: a quantitative analysis. Proc Natl Acad Sci U S A. 1990 Oct;87(19):7668–72.

25. de Smit MH, van Duin J. Control of translation by mRNA secondary structure in Escherichia coli. A quantitative analysis of literature data. J Mol Biol. 1994 Nov 25;244(2):144–50.

26. Evfratov SA, Osterman IA, Komarova ES, Pogorelskaya AM, Rubtsova MP, Zatsepin TS, et al. Application of sorting and next generation sequencing to study 5’-UTR influence on translation efficiency in Escherichia coli. Nucleic Acids Res. 2017 Apr 7;45(6):3487–502.

27. Yarchuk O, Jacques N, Guillerez J, Dreyfus M. Interdependence of translation, transcription and mRNA degradation in the lacZ gene. J Mol Biol. 1992 Aug 5;226(3):581–96.

28. Mearls EB, Jackter J, Colquhoun JM, Farmer V, Matthews AJ, Murphy LS, et al. Transcription and translation of the sigG gene is tuned for proper execution of the switch from early to late gene expression in the developing Bacillus subtilis spore. PLoS Genet. 2018 Apr 27;14(4):e1007350.

29. Shen X, Song S, Li C, Zhang J. Synonymous mutations in representative yeast genes are mostly strongly non-neutral. Nature. 2022 Jun 8;

30. Laabei M, Recker M, Rudkin JK, Aldeljawi M, Gulay Z, Sloan TJ, et al. Predicting the virulence of MRSA from its genome sequence. Genome Res. 2014 May;24(5):839–49.

31. Read TD, Massey RC. Characterizing the genetic basis of bacterial phenotypes using genome-wide association studies: a new direction for bacteriology. Genome Med. 2014;6(11):109.

32. Priest NK, Rudkin JK, Feil EJ, van den Elsen JMH, Cheung A, Peacock SJ, et al. From genotype to phenotype: can systems biology be used to predict Staphylococcus aureus virulence? Nat Rev Microbiol. 2012 Nov;10(11):791–7.

33. Denamur E, Condamine B, Esposito-Farèse M, Royer G, Clermont O, Laouenan C, et al. Genome wide association study of Escherichia coli bloodstream infection isolates identifies genetic determinants for the portal of entry but not fatal outcome. PLoS Genet. 2022 Mar;18(3):e1010112.

34. Young BC, Wu CH, Charlesworth J, Earle S, Price JR, Gordon NC, et al. Antimicrobial resistance determinants are associated with Staphylococcus aureus bacteraemia and adaptation to the healthcare environment: a bacterial genome-wide association study. Microb Genom. 2021 Nov;7(11).

35. Mozos IR de los, Vergara-Irigaray M, Segura V, Villanueva M, Bitarte N, Saramago M, et al. Base Pairing Interaction between 5’- and 3’-UTRs Controls icaR mRNA Translation in Staphylococcus aureus. PLOS Genetics. 2013 Dec 19;9(12):e1004001.

36. Dalla Serra M, Coraiola M, Viero G, Comai M, Potrich C, Ferreras M, et al. Staphylococcus aureus Bicomponent γ-Hemolysins, HlgA, HlgB, and HlgC, Can Form Mixed Pores Containing All Components. J Chem Inf Model. 2005 Nov 1;45(6):1539–45.

37. Balasubramanian D, Ohneck EA, Chapman J, Weiss A, Kim MK, Reyes-Robles T, et al. Staphylococcus aureus Coordinates Leukocidin Expression and Pathogenesis by Sensing Metabolic Fluxes via RpiRc. mBio [Internet]. 2016 Jun 21 [cited 2020 Apr 21];7(3). Available from: https://www.ncbi.nlm.nih.gov/pmc/articles/PMC4916384/

38. Bronner S, Stoessel P, Gravet A, Monteil H, Prévost G. Variable Expressions of Staphylococcus aureus Bicomponent Leucotoxins Semiquantified by Competitive Reverse Transcription-PCR. Appl Environ Microbiol. 2000 Sep 1;66(9):3931–8.

39. Dunman PM, Murphy E, Haney S, Palacios D, Tucker-Kellogg G, Wu S, et al. Transcription Profiling-Based Identification ofStaphylococcus aureus Genes Regulated by the agrand/or sarA Loci. Journal of Bacteriology. 2001 Dec 15;183(24):7341–53.

40. Liang X, Yu C, Sun J, Liu H, Landwehr C, Holmes D, et al. Inactivation of a two-component signal transduction system, SaeRS, eliminates adherence and attenuates virulence of Staphylococcus aureus. Infect Immun. 2006 Aug;74(8):4655–65.

41. Flack CE, Zurek OW, Meishery DD, Pallister KB, Malone CL, Horswill AR, et al. Differential regulation of staphylococcal virulence by the sensor kinase SaeS in response to neutrophil-derived stimuli. Proc Natl Acad Sci U S A. 2014 May 13;111(19):E2037–45.

42. Voyich JM, Vuong C, DeWald M, Nygaard TK, Kocianova S, Griffith S, et al. The SaeR/S Gene Regulatory System is Essential for Innate Immune Evasion by Staphylococcus aureus. J Infect Dis. 2009 Jun 1;199(11):1698–706.

43. Yamazaki K, Kato F, Kamio Y, Kaneko J. Expression of γ-hemolysin regulated by sae in Staphylococcus aureus strain Smith 5R. FEMS Microbiol Lett. 2006 Jun 1;259(2):174–80.

44. Nygaard TK, Pallister KB, Ruzevich P, Griffith S, Vuong C, Voyich JM. SaeR binds a consensus sequence within virulence gene promoters to advance USA300 pathogenesis. J Infect Dis. 2010 Jan 15;201(2):241–54.

45. Saïd-Salim B, Dunman PM, McAleese FM, Macapagal D, Murphy E, McNamara PJ, et al. Global Regulation of Staphylococcus aureus Genes by Rot. Journal of Bacteriology. 2003 Jan 15;185(2):610–9.

46. Venkatasubramaniam A, Kanipakala T, Ganjbaksh N, Mehr R, Mukherjee I, Krishnan S, et al. A Critical Role for HlgA in Staphylococcus aureus Pathogenesis Revealed by A Switch in the SaeRS Two-Component Regulatory System. Toxins. 2018 Sep;10(9):377.

47. Sun F, Li C, Jeong D, Sohn C, He C, Bae T. In the Staphylococcus aureus Two-Component System sae, the Response Regulator SaeR Binds to a Direct Repeat Sequence and DNA Binding Requires Phosphorylation by the Sensor Kinase SaeS. J Bacteriol. 2010 Apr;192(8):2111–27.

48. Yamashita K, Kawai Y, Tanaka Y, Hirano N, Kaneko J, Tomita N, et al. Crystal structure of the octameric pore of staphylococcal γ-hemolysin reveals the β-barrel pore formation mechanism by two components. Proc Natl Acad Sci U S A. 2011 Oct 18;108(42):17314–9.

49. Gu T, Zhao S, Pi Y, Chen W, Chen C, Liu Q, et al. Highly efficient base editing in Staphylococcus aureus using an engineered CRISPR RNA-guided cytidine deaminase. Chem Sci. 2018 Mar 28;9(12):3248–53.

50. Boisset S, Geissmann T, Huntzinger E, Fechter P, Bendridi N, Possedko M, et al. Staphylococcus aureus RNAIII coordinately represses the synthesis of virulence factors and the transcription regulator Rot by an antisense mechanism. Genes Dev. 2007 Jun 1;21(11):1353–66.

51. Hodille E, Alekseeva L, Berkova N, Serrier A, Badiou C, Gilquin B, et al. Staphylococcal Enterotoxin O Exhibits Cell Cycle Modulating Activity. Front Microbiol. 2016;7:441.

52. Desgranges E, Barrientos L, Herrgott L, Marzi S, Toledo-Arana A, Moreau K, et al. The 3’UTR-derived sRNA RsaG coordinates redox homeostasis and metabolism adaptation in response to glucose-6-phosphate uptake in Staphylococcus aureus. Molecular Microbiology. 2022;117(1):193–214.

53. Lalaouna D, Baude J, Wu Z, Tomasini A, Chicher J, Marzi S, et al. RsaC sRNA modulates the oxidative stress response of Staphylococcus aureus during manganese starvation. Nucleic Acids Res. 2019 Oct 10;47(18):9871–87.

54. Wolz C, Pöhlmann-Dietze P, Steinhuber A, Chien YT, Manna A, Van Wamel W, et al. Agr-independent regulation of fibronectin-binding protein(s) by the regulatory locus sar in Staphylococcus aureus. Molecular Microbiology. 2000 Apr 1;36(1):230–43.

55. Nonfoux L, Chiaruzzi M, Badiou C, Baude J, Tristan A, Thioulouse J, et al. Impact of Currently Marketed Tampons and Menstrual Cups on Staphylococcus aureus Growth and Toxic Shock Syndrome Toxin 1 Production In Vitro. Appl Environ Microbiol. 2018 May 31;84(12):e00351–18.

56. Geissmann T, Chevalier C, Cros MJ, Boisset S, Fechter P, Noirot C, et al. A search for small noncoding RNAs in Staphylococcus aureus reveals a conserved sequence motif for regulation. Nucleic Acids Research. 2009 Nov 1;37(21):7239–57.

57. Khusainov I, Vicens Q, Ayupov R, Usachev K, Myasnikov A, Simonetti A, et al. Structures and dynamics of hibernating ribosomes from Staphylococcus aureus mediated by intermolecular interactions of HPF. EMBO J. 2017 Jul 14;36(14):2073–87.

58. Arnaud M, Chastanet A, Débarbouillé M. New vector for efficient allelic replacement in naturally nontransformable, low-GC-content, gram-positive bacteria. Appl Environ Microbiol. 2004 Nov;70(11):6887–91.

59. Horinouchi S, Weisblum B. Nucleotide sequence and functional map of pC194, a plasmid that specifies inducible chloramphenicol resistance. J Bacteriol. 1982 May;150(2):815–25.

60. Nair D, Memmi G, Hernandez D, Bard J, Beaume M, Gill S, et al. Whole-genome sequencing of Staphylococcus aureus strain RN4220, a key laboratory strain used in virulence research, identifies mutations that affect not only virulence factors but also the fitness of the strain. J Bacteriol. 2011 May;193(9):2332–5.

61. Diep BA, Palazzolo-Ballance AM, Tattevin P, Basuino L, Braughton KR, Whitney AR, et al. Contribution of Panton-Valentine leukocidin in community-associated methicillin-resistant Staphylococcus aureus pathogenesis. PLoS One. 2008 Sep 12;3(9):e3198.

62. Perret M, Badiou C, Lina G, Burbaud S, Benito Y, Bes M, et al. Cross-talk between Staphylococcus aureus leukocidins-intoxicated macrophages and lung epithelial cells triggers chemokine secretion in an inflammasome-dependent manner. Cell Microbiol. 2012 Jul;14(7):1019–36.

63. Vandenesch F, Naimi T, Enright MC, Lina G, Nimmo GR, Heffernan H, et al. Community-acquired methicillin-resistant Staphylococcus aureus carrying Panton-Valentine leukocidin genes: worldwide emergence. Emerging Infect Dis. 2003 Aug;9(8):978–84.

64. Herbert S, Ziebandt AK, Ohlsen K, Schäfer T, Hecker M, Albrecht D, et al. Repair of global regulators in Staphylococcus aureus 8325 and comparative analysis with other clinical isolates. Infect Immun. 2010 Jun;78(6):2877–89.

65. Marincola G, Schäfer T, Behler J, Bernhardt J, Ohlsen K, Goerke C, et al. RNase Y of Staphylococcus aureus and its role in the activation of virulence genes. Mol Microbiol. 2012 Sep;85(5):817–32.

66. Redder P, Linder P. New Range of Vectors with a Stringent 5-Fluoroorotic Acid-Based Counterselection System for Generating Mutants by Allelic Replacement in Staphylococcus aureus. Appl Environ Microbiol. 2012 Jun;78(11):3846–54.

67. Diep BA, Gill SR, Chang RF, Phan TH, Chen JH, Davidson MG, et al. Complete genome sequence of USA300, an epidemic clone of community-acquired meticillin-resistant Staphylococcus aureus. Lancet. 2006 Mar 4;367(9512):731–9.

68. Fey PD, Endres JL, Yajjala VK, Widhelm TJ, Boissy RJ, Bose JL, et al. A genetic resource for rapid and comprehensive phenotype screening of nonessential Staphylococcus aureus genes. mBio. 2013 Feb 12;4(1):e00537–00512.

69. Charpentier E, Anton AI, Barry P, Alfonso B, Fang Y, Novick RP. Novel cassette-based shuttle vector system for gram-positive bacteria. Appl Environ Microbiol. 2004 Oct;70(10):6076–85.

70. Gruber AR, Lorenz R, Bernhart SH, Neuböck R, Hofacker IL. The Vienna RNA websuite. Nucleic Acids Res. 2008 Jul 1;36(Web Server issue):W70–74.

